# Long-term protection of rhesus macaques from Zika virus reinfection

**DOI:** 10.1101/712281

**Authors:** Gage K. Moreno, Christina M. Newman, Michelle R. Koenig, Mariel S. Mohns, Andrea M. Weiler, Sierra Rybarczyk, Logan J. Vosler, Nicholas Pomplun, Nancy Schultz-Darken, Eva Rakasz, Dawn M. Dudley, Thomas C. Friedrich, David H. O’Connor

**Affiliations:** Department of Pathology and Laboratory Medicine, University of Wisconsin-Madison, Madison, WI, United States of America; Wisconsin National Primate Research Center, University of Wisconsin-Madison, Madison, WI, United States of America; Department of Pathobiological Sciences, University of Wisconsin-Madison, Madison, WI, United States of America

**Author notes:** These authors contributed equally to this work (Gage K. Moreno, Christina M. Newman).

## Abstract

By the end of the 2016 Zika virus (ZIKV) outbreak, it is estimated that there were up to 100 million infections in the Americas. In approximately one in seven infants born to mothers infected during pregnancy, ZIKV has been linked to microcephaly, developmental delays, or other congenital disorders collectively known as congenital Zika syndrome (CZS). Guillain-Barré syndrome (GBS) in ZIKV infected adults. It is a global health priority to develop a vaccine against ZIKV that elicits long-lasting immunity, however, the durability of immunity to ZIKV is unknown. Previous studies in mice and nonhuman primates have been crucial in vaccine development but have not defined the duration of immunity generated by ZIKV infection. In this study, we rechallenged five rhesus macaques with ZIKV two years after a primary ZIKV infection. We show that primary ZIKV infection generates high titers of neutralizing antibodies (nAbs) that protect from detectable plasma viremia following rechallenge and persist for at least 27 months. While additional longitudinal studies are necessary with longer time frames, this study establishes a new experimentally defined minimal length of protective ZIKV immunity.

**Author Summary:** ZIKV emerged as a vector-borne pathogen capable of causing illness in infected adults and congenital birth defects in infants born to mothers infected during pregnancy. Despite the drop in ZIKV cases since the 2015-16 epidemic, questions concerning the prevalence and longevity of protective immunity have left vulnerable communities fearful that they may become the center of next ZIKV outbreak. While pre-existing herd immunity in regions of past outbreaks may dampen the potential for future outbreaks to occur, we currently do not know the longevity of protective immunity to ZIKV after a person becomes infected. Here, we establish a new experimentally defined minimal length of protective ZIKV immunity. We show that five rhesus macaques initially infected with ZIKV two years prior to rechallenge elicit a durable immune response that protected from detectable plasma viremia. While this work establishes a new minimal length of protective immunity, additional studies are necessary to define the maximum length of protective immunity following ZIKV infection.

## Introduction

Zika virus (ZIKV) is a mosquito-borne flavivirus first isolated in 1947 in the Zika forest of Uganda [1]. ZIKV is contracted through a mosquito bite or sexual contact and can be passed from mother to fetus during pregnancy [2]. ZIKV infections are typically mild: characterized by low-grade fever, maculopapular rash, arthralgia, and conjunctivitis [3]. The recent outbreak of ZIKV in the Americas was associated with more severe disease. Guillain-Barré syndrome (GBS) has found in a small proportion of ZIKV-positive adults in the French Polynesian and American outbreaks [4, 5]. In addition, infants born to mothers infected with ZIKV during pregnancy have an approximately one in seven chance of having congenital Zika syndrome (CZS), a collection of fetal abnormalities and developmental disorders [6, 7]. A retrospective analysis of infants, now aged 7 - 32 months, born to mothers infected with ZIKV during pregnancy, has shown that one-third of those infants have below average neurodevelopment [8]. Currently, it is unknown if ZIKV infection in humans confers lifelong immunity, however it is suggested that waning immunity over time leads to more severe ZIKV in the future [9, 10]. It is thought that a single infection or vaccination that generates neutralizing antibody (nAb) titers above 1:10 PRNT_50_ (50% plaque reduction neutralization test) is enough to confer lifelong immunity in other flaviviruses, such as yellow fever virus (YFV) or dengue virus (DENV) [11, 12]. Antibodies to YFV and DENV have been detected at least 60 years and 35 years after infection, respectively [13–15]. Despite long-term immunity to YFV and DENV, challenges remain in eradication of these viruses due to evidence of circulation in sylvatic (forest) cycles in Asia and Africa, and urban human-mosquito cycles in South America [16–19]. ZIKV circulates as one serotype while DENV circulates as four antigenically distinct serotypes (DENV-1 through DENV-4) [20, 21]. No single serotype of DENV has been eradicated after an outbreak even though there is long-term population-level immunity [15]. Similarly, there is a highly effective YFV vaccine that induces lifelong immunity but hasn’t led to the eradication of YFV [14]. Little is known about ZIKV transmission cycles in the Americas, but looking at past flavivirus outbreaks, herd immunity and vector control alone will likely not be enough to eradicate the virus. Determining the longevity of immunity elicited from previous natural ZIKV infection will be important for assessing the risk of reinfection in future outbreaks, as well as for estimating the number of people who are at-risk of future ZIKV infections.

The current understanding of ZIKV immunity in humans is derived primarily from vaccine clinical trials that are in progress [22]. Recipients of a prM+E consensus DNA vaccine (GLS-5700) developed nAb titers ranging from 1:18 to 1:317 fourteen weeks after the first vaccination. Adoptive transfer of this week fourteen serum provided protection in 92% of mice challenged with a Puerto Rican isolate of ZIKV [23]. Similar work with three purified inactivated virus (PIV) vaccines found that 92% of recipients generated nAb titers greater than 1:100 which corresponds to protection from viremia in mice that were challenged after adoptive antibody transfer of day 57 purified IgG [24]. ZIKV vaccine studies have shown that a range of nAb titers can protect from viremia. However, these studies did not look past 14 weeks to examine the duration of protection of vaccine-derived nAbs.

Several animal models, including mice and rhesus macaques, have been used to explore vaccine protection against experimental ZIKV exposure [25]. Immunocompromised mouse models have been used to advance the knowledge of ZIKV due to their cost effectiveness and availability. Published ZIKV challenge intervals in mice have varied from 3 to 30 weeks after vaccine administration and shown that nAb titers of EC50 (half maximal effective concentration) ranging from 1:75 to 1:100,000 were protective from viremia after ZIKV challenge [26–32]. While immunodeficient mouse models are readily accessible, ZIKV infection is typically lethal because these mice lack necessary components of the antiviral response. Immunodeficient mice provide information on efficacy and safety of therapeutic interventions but fail to provide information on natural virus pathogenicity [25, 33]. Non-human primates (NHP) have also been used in ZIKV studies because they more closely mimic human ZIKV infections [25]. For this reason, NHP studies have been used to assess the durability of immunity from 4 weeks to 1 year following vaccine administration [31, 32, 34, 35]. Looking at the longest duration between vaccination and challenge, Abbink et al. [32] conducted a study using rhesus macaques to demonstrate the protective efficacy of a ZIKV purified inactivated virus (PIV), DNA-M-Env, and RhAd52-M-Env vaccine candidates. At 52 weeks after administration, they challenged the macaques with Zika Virus/H.sapiens-tc/BRA/2015/Brazil-ZKV2015 (ZIKV-BR). The PIV and RhAd52-M-Env vaccines generated log_10_ MN50 (50% reduction of microneutralization, comparable to EC50 and PRNT50 values [36]) titers of 1.88 and 2.38 by week 4, but by week 8 after boost immunization log_10_ MN50 titers increased to and fluctuated between 3.71 and 2.42 through week 52, respectively. The DNA-M-Env vaccine generated log_10_ MN50 titers of 2.23 at week 8 after boost immunization and rapidly decline to 1.43 by week 14 but remained stable until week 52. These titers of neutralizing antibodies showed that the PIV, RhAd52-M-Env, and DNA-M-Env vaccines protected 75% (6 of 8), 100% (4 of 4), and 29% (2 of 7), respectively, of macaques after ZIKV-BR challenge [32].

Relying on vaccine studies is not sufficient to determine the durability of immunity because vaccine studies rely on booster vaccinations to increase the level of immunity [37]. Natural infections elicit a higher level of immunity that plateaus above the protective threshold thereby maintaining longer protection [38]. For this reason, rechallenge experiments are useful to understand the durability of immunity elicited by natural infection.

Rechallenge experiments using animal models have been used to examine immune responses following natural ZIKV infection, and to confirm that these infections protect from subsequent exposures [25]. Mouse models have been used to rechallenge 3 weeks after primary infection [39–41]. In rhesus macaques, challenge periods have varied from 6 weeks to 10 weeks after primary challenge [42–45]. We previously demonstrated that rhesus macaques were protected from homologous or heterologous rechallenge up to 10 weeks after primary infection [44, 45]. These studies have shown that following ZIKV rechallenge, CD8 T cell and CD4 T cell responses are weak, but there is an increase in natural killer (NK) cells and nAbs, which correspond to protection from disease but not sterilizing immunity [41, 42, 45]. While this is useful for establishing the potency of naturally elicited immunity, rapid rechallenge does not mimic the situation where someone infected in the 2015/2016 American outbreak is re-exposed years later in a subsequent outbreak.

Here we demonstrate a new minimum length of protection resulting from primary ZIKV infection using Indian-origin rhesus macaques from previously published studies [44, 46, 47].

## Results

### No detectable ZIKV in plasma following rechallenge

Five animals were infected with ZIKV as part of previously published studies [44–47]. Animal demographics, infection history, and naming scheme used for each animal are described in **Table 1**. Following these initial ZIKV infections, peak plasma vRNA load occurred between 2 and 6 days post infection (DPI) and ranged from 7.0 × 10^5^ to 2.8 × 10^4^ vRNA copies per ml [44–47]. In non-pregnant animals, vRNA load became undetectable by 10-14 DPI. In animals initially infected during pregnancy, vRNA load remained detectable for a longer period of time, ranging from 9-70 DPI [46]. To determine the durability of immunity following primary ZIKV infection, we subcutaneously rechallenged these 5 animals with 1 × 10^4^ PFU Zika virus/H.sapiens-tc/PUR/2015/PRVABC59 (ZIKV-PR) 19-27 months following initial ZIKV infection (**Table 1**). After rechallenge, ZIKV was measured in plasma by qRT-PCR daily on days 3-7 post rechallenge, and on days 10, 14, and 28 days post rechallenge (**Fig 1**). On all days throughout the study, plasma vRNA loads remained negative, indicating protection from plasma viremia after ZIKV rechallenge.

**Fig 1:**
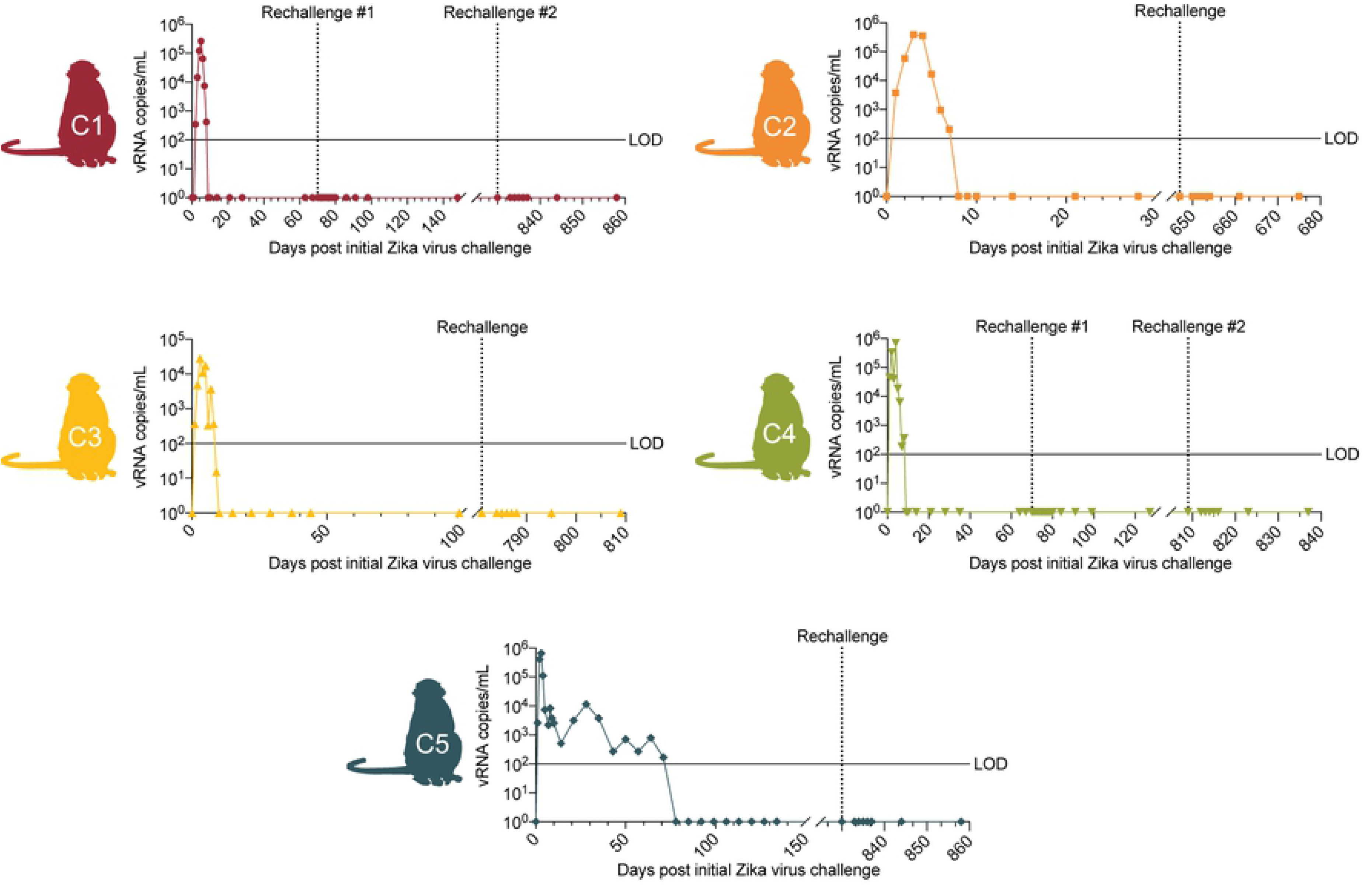
ZIKV viral loads of animals rechallenged with ZIKV-PR from day 0 of primary infection. Schematic of animals included in this study next to their plasma vRNA loads measured by qRT-PCR before and after rechallenge with ZIKV-PR as indicated by an arrow. The limit of detection of ZIKV RNA is 100 copies/mL as indicated by the solid line.

**Table 1:** Animal demographics.

| Animal | Public Identifier | Sex | Initial ZIKV Inoculation strain | Initial inoculation dose and route | Rechallenge ZIKV Inoculation strain | Rechallenge inoculation dose, route, and days post initial ZIKV infection (dpi) | Other infections and days post initial ZIKV infection (dpi) |
| --- | --- | --- | --- | --- | --- | --- | --- |
| C1 | 295022 | Female | Zika virus/R.macaqu<br>e-<br>tc/UGA/1947/M<br>R766- 3329 | 10E4 PFU,<br>subcutaneo<br>us | Zika virus/H.sapiens-tc/ FRA/2013/<br>FrenchPolynesia-01_v1c | 10E4 PFU,<br>subcutaneous, 70 dpi | SHIV, 347 dpi |
|  |  |  |  | 10E4 PFU,<br>subcutaneo<br>us | Zika virus/H.sapiens-tc/<br>PUR/2015/PRVABC5<br>9 | 10E4 PFU,<br>subcutaneous, 858 dpi |  |
| C2 | 321142 | Female | Zika virus/H.sapiens-tc/ FRA/2013/<br>FrenchPolynesi<br>a- 01_v1c1 | 10E6 PFU,<br>subcutaneo<br>us | Zika virus/H.sapiens-tc/<br>PUR/2015/PRVABC5<br>9 | 10E4 PFU,<br>subcutaneous, 675 dpi | DENV3, 566 dpi |
| C3 | 598248 | Female | Zika virus/H.sapiens-tc/ FRA/2013/<br>FrenchPolynesi<br>a- 01_v1c1 | 10E4 PFU,<br>subcutaneo<br>us | Zika virus/H.sapiens-tc/<br>PUR/2015/PRVABC5<br>9 | 10E4 PFU,<br>subcutaneous, 809 dpi | None |
| C4 | 610107 | Female | Zika virus/H.sapiens-tc/ FRA/2013/<br>FrenchPolynesi<br>a- 01_v1c1 | 10E6 PFU,<br>subcutaneo<br>us | Zika virus/H.sapiens-tc/ FRA/2013/<br>FrenchPolynesia-01_v1c1 | 10E6 PFU,<br>subcutaneous, 70 dpi | SHIV, 326 dpi |
|  |  |  |  | 10E4 PFU,<br>subcutaneo<br>us | Zika virus/H.sapiens-tc/<br>PUR/2015/PRVABC5<br>9 | 10E4 PFU,<br>subcutaneous, 837 dpi |  |
| C5 | 827577 | Female | Zika virus/H.sapiens-tc/ FRA/2013/ FrenchPolynesia- 01_v1c1 | 10E4 PFU, subcutaneous | Zika virus/H.sapiens-tc/ PUR/2015/PRVABC59 | 10E4 PFU, subcutaneous, 858 dpi | None |

Public animal identifiers are used in studies on the Zika Open Research Portal
(https://zika.labkey.com/project/OConnor/begin.view?).

### Immune parameters do not change significantly following ZIKV rechallenge

In order to characterize the immune responses following a long-term ZIKV rechallenge, we assessed immune activation by measuring Ki-67 expression, as a marker of activation, in natural killer (NK) cells, CD4, CD8, and CD8 effector memory (T_EM_) T cells by flow cytometry (**S1 Fig**). The percentage of activated cells per microliter of blood are presented as fold-change from the number of activated cells found 3 days before rechallenge (baseline) to determine fold-change in activation from baseline (**S1 Fig**). Repeated measures ANOVA showed that there were no significant changes in the number of activated CD4, CD8, and CD8 T_EM_ T cell populations over time. In contrast, there was significant variation from baseline measurements in activated (Ki67+) NK cell populations (F(2.263, 9.051) = 10.36, p = 0.0039), but further analysis by Dunnett’s multiple comparisons test showed no significant change in NK cell populations when comparing number of activated NK cells each day after rechallenge to the number 3 days before rechallenge.

Serum nAb responses were measured using a flow cytometry-based ZIKV neutralization assay. Prior to rechallenge, all animals had high EC50 (50% effective concentration) values (**Fig 2A**), that were greater than the 1:10 PRNT50 values recommended for other flavivirus vaccines [11, 12]. Using a one-way paired t-test, we compared EC50 values on day 0 to EC50 values on day 14 and day 28 (**Fig 2B**). All animals showed increased nAb titers following rechallenge as noted by a greater EC50 value on day 14 compared to day 0 (p = 0.0025, t=5.604, df=4) and on day 28 (p =0.0015, t=6.446, df=4). The increase in nAbs shows that initial infection may fail to provide sterilizing immunity. Alternatively, it is possible that the antigen present in the virus preparation was sufficient to boost pre-existing antibody responses. However, the combination of immunophenotyping data, neutralization assay data, and persistent negative plasma ZIKV vRNA loads following rechallenge suggest that primary ZIKV infection is sufficient to protect from detectable viremia in the blood.

**Fig 2:**
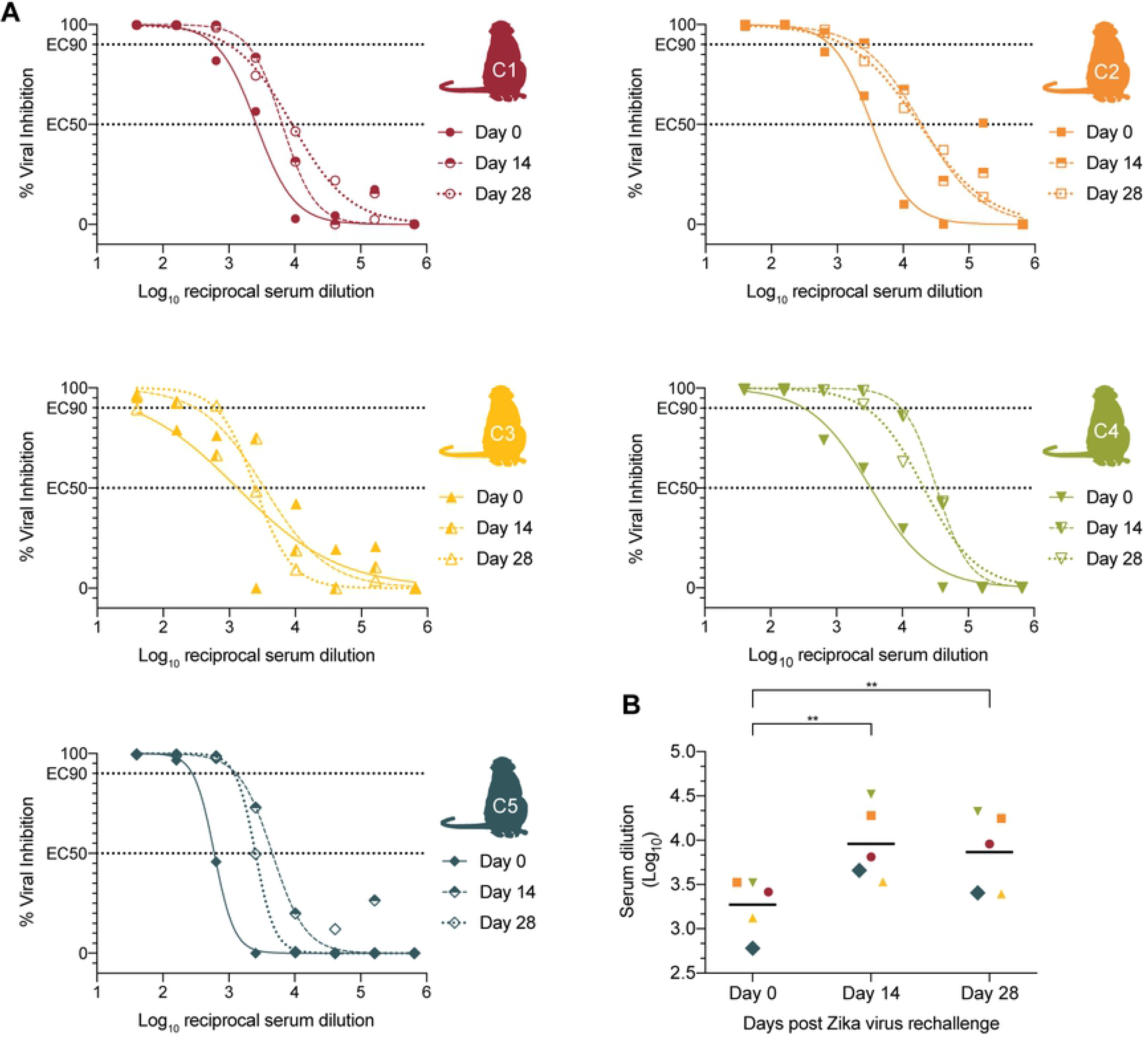
Neutralizing antibody curves. ZIKV neutralizing antibody titers were measured on days 0, 14, and 28. (A) The % viral inhibition corresponding to that log_10_ reciprocal serum dilution are plotted and fit with a non-linear regression line in order to estimate EC50 and EC90 values. (B) The log_10_ serum dilutions corresponding to EC50 nAb titers are plotted for days 0, 14, and 28. Increases in titers were measured by one-way paired t-test. Significance is indicated by ** (p-value <0.005).

### Clinical evaluation of ZIKV rechallenged macaques

After initial ZIKV challenge and rechallenge, all five animals were monitored daily for signs of disease (for example, inappetence, dehydration, diarrhea, injury, or psychological abnormalities) and had complete blood counts (CBCs) evaluated. Reference intervals (RIs) developed for Wisconsin National Primate Research Center (WNPRC) animals for species, sex, and age were used to evaluate results [44].

After primary ZIKV infection, animal C1 presented a rash at the injection site. No other clinical abnormalities were noted. Complete blood counts (CBCs) were evaluated for all five animals three days prior to challenge, daily on days 0 through 10, 14, and again on days 21 and 28 DPI. Mild monocytosis with levels exceeding the RI (0-3% of white blood cells (WBC)) was reported for all animals before initial infection and persisted throughout the duration of the study [44–47]. Animal C1 had lymphocyte levels dip below the RI (20-80% WBC) on days 4 and 7 but rebounded back to within the normal range by day 8. Animal C5 had high platelet counts exceeding the RI (229-524 thousands of platelets/uL) present at day −3 and persisted to be above the RI throughout the study period [44–47].

After rechallenge, animals C1 and C4 developed inappetence 6 days post rechallenge. Animal C1 also developed diarrhea at 6 days post rechallenge. All animals were relocated to new housing 1 day prior to rechallenge which may have contributed to inappetence and diarrhea. No other clinical abnormalities were noted (**S1 Table**). CBC tests were evaluated for all five animals 3 days prior to, on the day of rechallenge, and on days 3-7, 10, 14, and 28 post rechallenge (**S2 Fig**). After rechallenge, all animals displayed persistent mild monocytosis with levels exceeding the RI across all days that CBCs were evaluated. This persistence of monocytosis is consistent with that observed following primary ZIKV infection [44–47]. The stress of frequent blood draws, sedation, experimental infections, and living in captivity have all been associated with an increased risk of monocytosis and lymphopenia potentially explaining these findings in rechallenged animals [48–50].

## Discussion

Since ZIKV could re-emerge in areas affected by the 2015 ZIKV outbreak, it is important to evaluate the durability of ZIKV immunity following initial infection in order to understand the risk of re-infection, particularly among women. This study shows that initial ZIKV infection provides apparently protective immunity at least 27 months after initial ZIKV infection. Primary ZIKV infection generated nAb titers above the 1:10 threshold deemed protective against re-infection for other flaviviruses (**Fig 2**) [11, 12].

These results showing protection from long term rechallenge have important implications for vaccine design and testing. Protection from ZIKV infection following vaccination is thought to require lasting MN50 log_10_ titers of 2.0 - 2.1, equivalent to log_10_ EC50 titers. [32]. In our study, prior to rechallenge, EC50 nAb titers were well above the log_10_ 2.0 threshold that Abbink et al., determined to be protective from ZIKV infection [32]. Does the immunity observed in these animals provide sterilizing protection from reinfection? While we did not detect any ZIKV RNA in the plasma following rechallenge, nAb titers increased modestly. This suggests that ZIKV may replicate at levels below the limit of detection in blood or in sanctuary tissues that are sequestered from the blood. Conversely, it is possible that antigen present in the challenge virus itself was sufficient to stimulate an expansion of pre-existing nAb responses. Others who have encountered similar results have defined empirical thresholds for sterilizing immunity. For example, Kirkpatrick et al. similarly observed a boost in nAb titers in DENV-vaccinated animals following DENV challenge [51]. They defined an increase of less than four-fold nAb titer as sterilizing immunity and a ≥ four-fold increase in nAb titer as non-sterilizing immunity. In our study, two animals, C1 and C3, exhibited 2.5x increases in nAb titer by 14 days post exposure. However, animals C2, C4, and C5 exhibited a 5.7x, 10.0x, and 7.6x increase, respectively, in nAb titer by 14 days post exposure.

Two animals used in our rechallenge study, C3 and C5, were previously part of a published study that assessed fetal outcomes following ZIKV infection in utero [46]. This study concluded that similar to human pregnancy, maternal-fetal transmission is highly efficient with variable vRNA levels detected in all fetuses assessed. Despite a small sample size in our rechallenge study (n = 2 animals initially infected during pregnancy), we show that these animals develop immune responses similar to non-pregnant animals. Following primary ZIKV infection during pregnancy in 2016, animal C5 has gone on to give birth to two apparently healthy infants (in 2017 and 2018) and C3 has gone on to give birth to one healthy infant (2019). While this study did not look at the impacts of a ZIKV rechallenge during a second pregnancy, we show that subsequent infants born to mothers who were previously infected with ZIKV are in qualitatively good health. Future studies will need to determine if immunity elicited by a primary infection is sufficient to protect a subsequent pregnancy from the effects of a ZIKV rechallenge during pregnancy. Additionally, it will be important to characterize the extent of ZIKV-specific antibody responses in infants exposed to ZIKV *in utero*.

In summary, our study establishes a new experimentally defined minimal length of protective immunity against ZIKV of up to 27 months. Future longitudinal studies will need to assess ZIKV immunity of previously infected individuals at longer time frames than we have in order to continue to define the maximum length of immunity elicited by natural ZIKV infection. Establishing the true maximum length of ZIKV protective immunity will be critical for understanding outbreak dynamics, the future transmission potential of ZIKV, and influence future vaccination strategies.

## Materials and Methods

### Ethics Statement

All rhesus macaques in this study were cared for by the staff at the Wisconsin National Primate Research Center (WNPRC) in accordance with the regulations, guidelines, and recommendations outlined in the Animal Welfare Act, the Guide for the Care and Use of Laboratory Animals, and the Weatherall report [52–54]. The University of Wisconsin - Madison, College of Letters and Science and Vice Chancellor for Research and Graduate Education Centers Institutional Animal Care and Use Committee approved the nonhuman primate research covered under protocol number G005401-R01. The University of Wisconsin - Madison Institutional Biosafety Committee approved this work under protocol number B00000117.

All animals were housed in enclosures with required floor space and fed using a nutritional plan based on recommendations published by the National Research Council. Animals were fed a fixed formula, extruded dry diet with adequate carbohydrate, energy, fat, fiber, mineral, protein, and vitamin content. Macaque dry diets were supplemented with fruits, vegetables, and other edible objects (e.g., nuts, cereals, seed mixtures, yogurt, peanut butter, popcorn, marshmallows, etc.) to provide variety to the diet and to inspire species-specific behaviors such as foraging. To further promote psychological well-being, animals were provided with food enrichment, structural enrichment, and/or manipulanda. Environmental enrichment objects were selected to minimize chances of pathogen transmission from one animal to another and from animals to care staff. While on study, all animals were evaluated by trained animal care staff at least twice each day for signs of pain, distress, and illness by observing appetite, stool quality, activity level, physical condition. Animals exhibiting abnormal presentation for any of these clinical parameters were provided appropriate care by attending veterinarians. Prior to all minor/brief experimental procedures, macaques were anesthetized with an intramuscular dose of ketamine (10 ml kg^-1^) prior to virus inoculation and blood collection and monitored regularly until fully recovered from anesthesia. Per WNPRC standard operating procedures for animals assigned to protocols involving the experimental inoculation of an infectious pathogen, environmental enhancement included constant visual, auditory, and olfactory contact with conspecifics, the provision of feeding devices which inspire foraging behavior, the provision and rotation of novel manipulanda (e.g., Kong toys, nylabones, etc.), and enclosure furniture (i.e., perches, shelves).

### Virus Stocks

Zika virus/H.sapiens-tc/FRA/2013/FrenchPolynesia-01_v1c1 (ZIKV-FP) (GenBank accession: KJ776791) was obtained from Xavier de Lamballerie (European Virus Archive, Marseille, France) and passage history is described in Dudley et al. [44]. African lineage Zika virus (ZIKV-MR766), and Zika virus/H.sapiens-tc/PUR/2015/PRVABC59 (GenBank accession: KU501215) were provided by Brandy Russell (CDC, Ft. Collins, CO). ZIKV-MR766 passage history has been described in Aliota et al [45]. Prior to rechallenge, animal C1 was previously infected with 10^4^ PFU of ZIKV-MR766 as described by Aliota et al. [45]. Animal C4 was previously infected with 10^6^ PFU of ZIKV-FP as described by Dudley et al [44]. Animals C5 and C3 were previously infected with 10^4^ PFU of ZIKV-FP during their first and late second/early third trimesters of pregnancy, respectively, as described by Nguyen et al. [46]. Animal C2 was previously infected with 10^6^ PFU ZIKV-FP as previously described by Breitbach et al. [47]. Full infection histories are described in **Table 1**.

### Inoculations

Virus stocks were thawed and diluted in PBS to 1 × 10^4^ PFU Zika virus/H.sapiens-tc/PUR/2015/PRVABC59 (ZIKV-PR) for each challenge. Prepared virus stocks were then loaded into a 1 ml syringe that was kept on ice until administration. Animals were anesthetized as described above, and 1 ml of inocula was administered subcutaneously over the cranial dorsum [44]. Animals were monitored for adverse reactions and signs of disease by the veterinary and animal care staff of WNPRC.

### Quantification of ZIKV RNA

Plasma and PBMC were isolated from EDTA-anticoagulated whole blood by Ficoll density centrifugation at 1,860 x g for 30 min. Plasma was collected and centrifuged at 670 x g for an additional 8 min to remove remaining cells. Viral RNA was extracted from 300μL of plasma using the Viral Total Nucleic Acid Purification kit (Promega, Madison, WI, USA) on a Maxwell 48 RSC instrument (Promega, Madison, WI). The vRNA was then quantified using quantitative RT-PCR previously described in Dudley et al. [44]. Briefly, the qRT-PCR was performed using SuperScript III Platinum One-Step Quantitative RT-PCR System (Invitrogen, Carlsbad, CA, USA) on a LightCycler 480 or LightCyc instrument (Roche Diagnostics, Indianapolis, IN, USA). Primers and probes were used at final concentrations of 600 and 100 nM, respectively, with the addition of 150 ng random primers (Promega, Madison, WI). Cycling conditions were as follows: 37°C for 15 min, 50°C for 30 min, and 95°C for 2 min, followed by 50 cycles of 95°C for 15 sec and 60°C for 1 min. Viral concentration was calculated by interpolation onto an internal standard curve composed of seven 10-fold serial dilutions of a ZIKV RNA fragment based on ZIKV-FP.

The limit of quantification for this assay was determined using methods adapted from Cline et al [55]. Briefly, half-dilutions of ZIKV RNA were prepared to generate standard curves from 3 to 1,000,000 copies. The overall positivity for each half dilution was assessed by qRT-PCR. The limit of quantification for this assay was established at 100 copies/ml.

### Immunophenotyping

The number of activated/proliferating NK cells, CD4, CD8, and CD8 T_EM_ T cells were quantified as described previously [56]. Briefly, 100μL of EDTA-anticoagulated whole blood samples were incubated at room temperature for 15 minutes with an antibody master mix described by Dudley et al. [44]. Red blood cells were lysed using BD Pharm Lyse (BD BioSciences, San Jose, CA), washed twice in media, and fixed with 125μL of 2% paraformaldehyde for 15 minutes. After fixation, cells were washed and permeabilized using Bulk Permeabilization Reagent (Life Technologies, Madison, WI). While the permeabilizer was present, the cells were stained for 15 minutes with Ki-67 (clone B56, Alexa Fluor 647 conjugate). The cells were then washed twice in media and resuspended in 2% paraformaldehyde until they were run on a BD LSRII Flow Cytometer (BD BioSciences, San Jose, CA). Flow data were analyzed using Flowjo version 10.5.3.

### CBC tests

CBC tests were performed on EDTA-anticoagulated whole-blood samples using a SYsmex XS-1000i automated haematology analyzer (Sysmex Corporation, Kobe, Japan). CBC results were reported with species, age, and sex-specific reference ranges. When samples are outside the laboratory-defined criteria (increased total WBC counts, increased monocyte, eosinophil, and basophil percentages, decreased hemoglobin, hematocrit, and platelet values, and unreported differential values) manual slide evaluation was performed.

### Flow cytometry-based neutralization assay

Macaque serum samples were screened for ZIKV nAbs by a flow-based ZIKV neutralization assay adapted from Dowd et al [20]. Briefly, Vero cells (ATCC, CLR-1587) were plated at a concentration of 2.5 × 10^4^ cells (100μ/well) in a 96 well plate one and incubated at 37°C for 24 hours prior to ZIKV infection. Equal volumes of 1.9 × 10^2^ PFU/mL ZIKV-PR and 1:4 serial serum dilutions were incubated for 30 minutes at 37°C before infecting duplicate wells. The plate was incubated for 72 hours at 37°C. Following incubation, the plates were washed 5 times with 1 × PBS and treated with trypsin. After the cells separated from the plate, they were transferred to a cluster tube and stained with a LIVE/DEAD Fixable Near-IR Dead Cell Stain Kit (Life Technologies, Madison, W). The plate was incubated for 15 minutes at room temperature, 1 mL of R10 media was added, and centrifuged for 5 minutes at 530 × g. Media was aspirated and 125μL of 2% PFA was added to fix the cells. The plates were incubated for 15 minutes at room temperature, 1 mL of R10 media was added, and then they were centrifuged for 5 minutes at 530 x g. Media was aspirated and 100μL of Fix and Permeabilization Medium B (Life Technologies, Madison, W) was added to permeabilize the cell membrane. 2μl of a 1:10 dilution of the flavivirus-specific antibody, 4G2-A647 (Novus Biologicals, Centennial, CO, USA) stock was added to each well. The plate was incubated at room temperature for 15 minutes. 1 mL of R10 media was added, and the plates were centrifuged for 5 minutes at 530 × g. Media was aspirated and 125μL of 2% PFA was added to fix the cells. Infection was monitored by flow cytometry on the BD LSRII Flow Cytometer. Flow cytometry data was analyzed using Flowjo version 10.5.3. Neutralization curves were generated using GraphPad Prism v. 8.0.1 (San Diego, CA). The dilutions required to inhibit 50% and 90% of infection (EC50 and EC90 values) were calculated using a non-linear regression model.

### Statistical Analysis

Changes in immunophenotyping and CBC parameters from baseline were analyzed with repeated measures ANOVA, for the populations where ANOVA proved significant, we followed with Dunnett’s multiple comparisons test comparing each day after rechallenge back to 3 days before rechallenge.

The log_10_ serum dilutions corresponding to EC50 nAb titers are plotted for days 0, 14, and 28. Increases in titers were measured by one-way paired t-test. Significance is indicated by * (p-value < 0.05) and ** (p-value <0.005). All statistical analysis was performed in GraphPad Prism v. 8.0.1 (San Diego, CA).

## Data management

Complete datasets for these studies have been made publicly available in a manuscript-specific folder on the Zika Open Research Portal (https://go.wisc.edu/bn71k1). Authors declare that all other data for these study findings are available via this portal or through supplementary information files from this article.

## Acknowledgments

We thank the Veterinary Services, Colony Management, Scientific Protocol Implementation, and the Pathology Services staff at the Wisconsin National Primate Research Center (WNPRC) for their contributions to this study.

## Supporting information captions

**S1 Fig: Measurement of immune activation of animals rechallenged with ZIKV-PR.** (A) Schematic of animals included in this study. (B) Spread of Ki-67+ NK cells, (C) CD4 T cells, (D) CD8 T cells, and (E) CD8 effector memory (T_EM_) T cells were measured on - 3, 0, 3-7, 10, 14, and 28 days post rechallenge. Percentage of activated cells per μl of blood are presented as relative to the baseline value set to 100% on −3 days post rechallenge.

**S2 Fig: Complete blood count graphs of animals rechallenged with ZIKV-PR.** Schematic of animals included in this study. (B) Clinical hematology was performed by WNPRC on whole blood collected daily from macaques on days 3-7, as well as on days −3, 0, 10, 14, and 28 post-ZIKV rechallenge. Horizontal dotted lines indicate the range of normal reference intervals (RI). Four of five animals had lymphocyte levels dip below the RI (20-80% of WBC) at 3 days post exposure but rebounded to the levels within the normal range by 4 days post exposure. Mild monocytosis exceeding the RI (0-3% of WBC) was reported in animal C2 7 days prior to rechallenge. Animal C5 had platelet values exceeding the RI (229-524 ths/uL) ranging from 575-747 ths/uL present at day −3 and continuing until day 28. Thrombocytosis was also noted following the initial ZIKV challenge for this animal. No other abnormal CBC data were reported.

**S1 Table: Clinical remarks for each animal after rechallenge.** BARH = Bright, Alert, Responsive, and Hydrated. BCS = Body Condition Score.

